# Aging Remodels the Female Skeletal Muscle Proteome Toward Matrix, Chromatin and Lysosomal Programs that Track Loss of Physical Function

**DOI:** 10.64898/2026.11.06.764992

**Authors:** Mark A. Watson, Christina D. King, Taylor R. Valentino, Heather L. Gregory, John Stone, Jamie N. Justice, Bumsoo Ahn, Birgit Schilling

## Abstract

Loss of skeletal muscle strength and mobility is among the most consequential features of human aging, yet the molecular programs that accompany this decline in women remain poorly understood. Here, we performed deep comprehensive data-independent acquisition (DIA) proteomic profiling of vastus lateralis biopsies from 10 younger (24-30 years) and 10 older (70-84 years) healthy women enrolled through the NIH Common Fund Cellular Senescence Network (SenNet), with matched measurements of grip strength and gait speed. Of 2,391 quantified protein groups, 163 were significantly altered with age, and partial least squares-discriminant analysis completely separated the age groups (cross-validated AUC = 1.00). Older muscle showed increased extracellular matrix and vascular remodeling proteins (TIMP3, COL8A1, ELN, FBN1), nuclear, chromatin and RNA-binding proteins (HMGB1, HMGB2, histone H1.0, MATR3, NCL), and lysosomal and senescence-associated secretory phenotype (SASP)-associated proteins (CTSZ, CTSD, PSAP), together with reduced glycolytic and energetic proteins (GAPDH, ENO3, PHKA1, CYCS). Age effects were robust to adjust for body mass index (BMI). Integrating differential abundance, multivariate importance and participant-level consistency, we defined a 24-protein skeletal muscle aging signature whose score separated all younger from all elderly women. Combining grip strength and gait speed into a combined function score, we found that 23 of the 24 signature proteins correlated with physical function (FDR < 0.05), and the signature score correlated strongly with function (r = −0.79). The senescence-associated alarmins HMGB1 and HMGB2 emerged as lead proteins: HMGB1 showed the strongest association with function (r = −0.86), and both remained associated after adjustment for age group. Together, these data define a senotype proteomic program in aging female skeletal muscle and a molecular signature of muscle function that, after validation, may serve as a readout for interventions that aim to preserve physical performance in elderly women.

## Introduction

Maintaining skeletal muscle health is one of the major challenges for healthy aging and for human health more broadly. Sarcopenia, the age-related loss of skeletal muscle mass, strength and function, affects an estimated 10-27% of adults aged 60 years and older worldwide, depending on the diagnostic criteria applied (Petermann-Rocha et al., 2022). Sarcopenia is a key contributor to frailty, a state of increased vulnerability to stressors that predisposes older adults to falls, disability, hospitalization and death (Clegg et al., 2013), and frailty is more prevalent in older women than in older men (Collard et al., 2012). Measures of muscle function are among the strongest predictors of health outcomes in older adults: lower grip strength is associated with higher all-cause and cardiovascular mortality (Leong et al., 2015), and slower gait speed is associated with shorter survival (Studenski et al., 2011). As populations age worldwide, preserving muscle strength and mobility has therefore become a central goal of geroscience and public health.

One of the most striking phenotypes of human aging is the progressive decline of skeletal muscle strength and physical performance, which contributes to impaired mobility, frailty, falls and loss of independence (Cruz-Jentoft et al., 2019; Studenski et al., 2014). Age-related loss of muscle mass and function, termed sarcopenia, was long regarded as part of normal aging, but was recognized with an ICD-10 diagnosis code in 2016 (Anker et al., 2016). Notably, the decline in strength exceeds what would be expected from the loss of muscle mass alone, suggesting that the intrinsic quality and contractile capacity of aging muscle tissue is altered (Zane et al., 2017; Gonzalez-Freire et al., 2018). Skeletal muscle health and homeostasis is increasingly recognized as arising from the integration of multiple biological processes, including metabolism and bioenergetics, proteostasis, genomic regulation, excitability, tissue structure, regeneration and intercellular cross-talk (Vainshtein et al., 2026). The molecular mechanisms that disrupt these interconnected processes within aging populations remain incompletely understood, and no treatment is currently available to prevent or delay progressive decline of muscle health.

Proteomic technologies offer a direct view of the functional effector molecules of muscle tissue. Previous proteomic studies of human skeletal muscle from the Baltimore Longitudinal Study of Aging (BLSA) and the Genetic and Epigenetic Signatures of Translational Aging Laboratory Testing (GESTALT) study revealed that older muscle is characterized by lower abundance of ribosomal and energetic proteins, including those of glycolysis, the tricarboxylic acid cycle and mitochondrial respiration, and by higher abundance of proteins involved in innate and adaptive immunity, proteostasis and alternative splicing (Ubaida-Mohien et al., 2019a). Physical activity was associated with partial preservation of the youthful muscle proteome (Ubaida-Mohien et al., 2019b), and single-fiber proteomics further demonstrated fiber type-specific features of muscle aging (Murgia et al., 2017). Structural and proteomic changes have also been described in muscle of physically active older men (Brocca et al., 2017). However, most of these studies included predominantly male participants or combined both sexes, and the proteomic remodeling of aging skeletal muscle specifically in women, and its relationship to clinical measures of muscle function, has received comparatively little attention.

Cellular senescence is a state of irreversible cell cycle arrest accompanied by the secretion of pro-inflammatory cytokines, chemokines, growth factors, proteases and ECM-modifying factors, collectively termed the senescence-associated secretory phenotype (SASP) (Coppé et al., 2008; Coppé et al., 2010; Gorgoulis et al., 2019; Basisty et al., 2020). Senescent cells accumulate with age in multiple tissues, are considered a hallmark of aging (López-Otín et al., 2023), and have been characterized in aging human skeletal muscle (Zhang et al., 2022). As there are no universal markers of senescence, the NIH Common Fund Cellular Senescence Network (SenNet) was established to map senescent cells (Suryadevara et al., 2024; Gurkar et al., 2023; Suryadevara et al., 2026) and specifically their molecular phenotypes, or <u>senotypes</u> (Schafer et al., 2026), across human tissues and the lifespan. Within SenNet, our Tissue Mapping Center has applied deep comprehensive proteomic profiling to human tissues, including the postmenopausal ovary, where we recently described dynamic matrix and fibroinflammatory remodeling in the decades after menopause (Watson et al., 2026).

Here, we extended this approach to skeletal muscle and performed deep comprehensive DIA proteomic profiling of vastus lateralis biopsies from younger and older healthy women with matched measurements of grip strength and gait speed. Combining differential abundance analysis, multivariate modeling and clinical correlation, we identified a coordinated program of matrix, nuclear and lysosomal remodeling with metabolic decline in older muscle, defined a 24-protein skeletal muscle aging signature, termed Buck SENmuscle 1.0, and show that this signature, led by HMGB1 and HMGB2, tracks a combined measure of physical function. Linking a defined tissue proteomic signature directly to clinically measurable functional metrics in the same individuals has rarely been achieved. Buck SENmuscle 1.0 therefore provides a candidate molecular readout of muscle function in aging women that can be tested in other cohorts and used to evaluate interventions aimed at preserving physical performance.

## Results

### Deep comprehensive proteomic profiling of aging female skeletal muscle

To characterize the effects of aging on the female skeletal muscle proteome we used in depth proteomic technologies, such as data-independent acquisitions mass spectrometry (DIA-MS). We analyzed muscle biopsies (vastus lateralis) from 20 healthy women enrolled through the SenNet Buck Tissue Mapping Center and Wake Forest University, comprising 10 younger (24-30 years, mean 26.6 ± 2.5 years) and 10 older participants (70-84 years, mean 76.8 ± 4.7 years) (Figure 1A, B; Figure S1; Table S1). Elderly women showed markedly lower grip strength (20.2 ± 5.1 vs. 30.8 ± 4.0 kg, p = 6.7 x 10^-5^) and slower gait speed (1.02 ± 0.14 vs. 1.17 ± 0.13, p = 0.020), whereas body mass index (BMI) was modestly lower in the older group (24.5 ± 4.7 vs. 28.4 ± 3.7 kg/m^2^, p = 0.054) (Figure 1B; Table S2). Muscle lysates were digested using S-Trap mini columns and analyzed by ‘parallel accumulation-serial fragmentation combined with data-independent acquisition’ (dia-PASEF) on a timsTOF HT mass spectrometer, and data were processed in Spectronaut using a library-free directDIA workflow (Figure 1A). In total, 2,770 protein groups were identified, of which 2,391 protein groups with at least two unique peptides were quantified across participants (Table S3). Quantification was highly complete (more than 2,385 protein groups per participant, < 0.1% missing values), with median coefficients of variation of 29% and 26% within the younger and older groups, respectively, and sample-to-sample Pearson correlations of 0.89 or higher (Figure S2).

**Figure 1.**
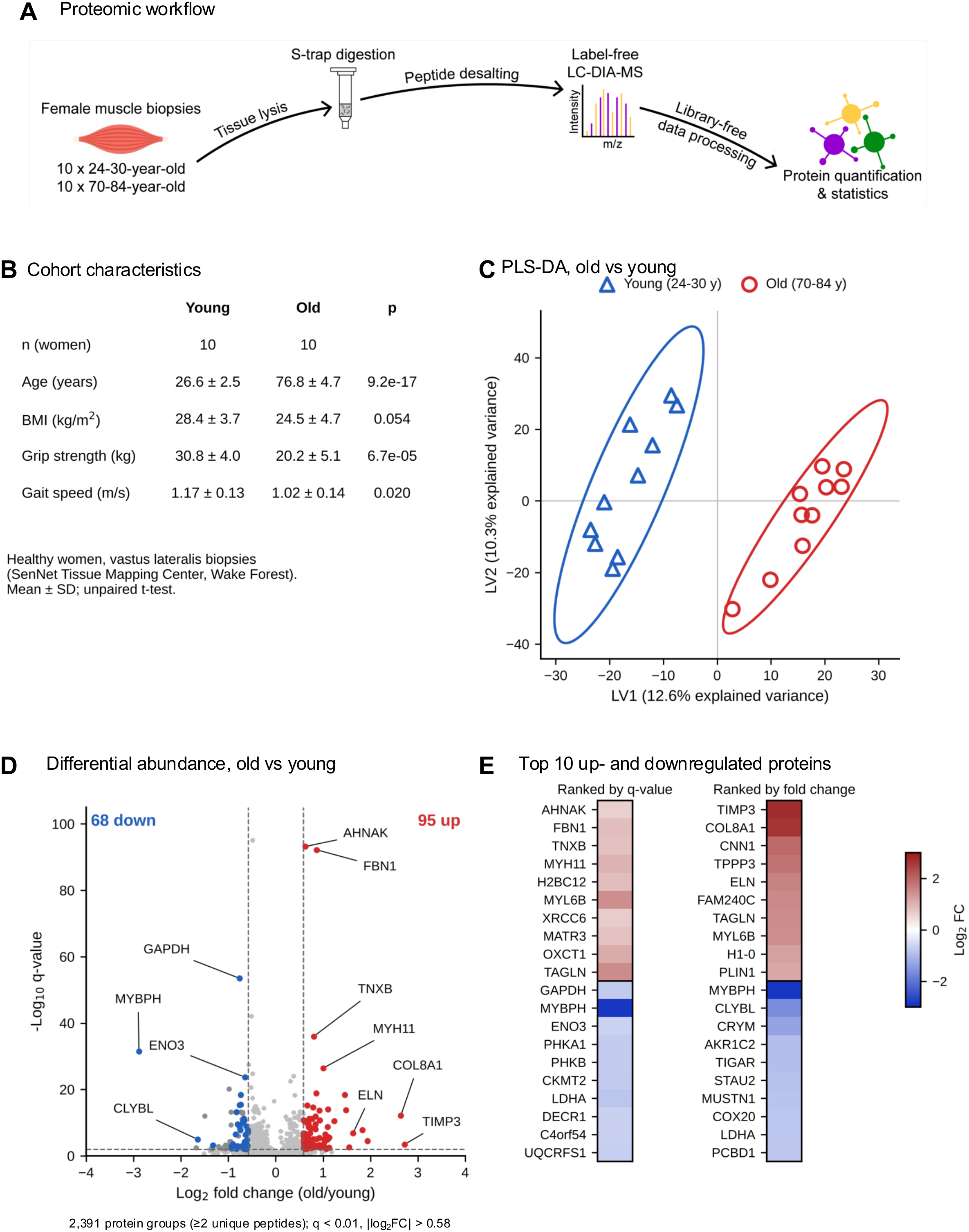
Deep comprehensive DIA proteomic profiling of aging female skeletal muscle. (A) Proteomic workflow. Vastus lateralis biopsies were lysed, digested on S-Trap columns, desalted and analyzed by dia-PASEF, followed by library-free directDIA data processing in Spectronaut. (B) Cohort characteristics (mean ± SD; unpaired t-test). Gait speed units to be confirmed. (C) PLS-DA score plot of 2,391 quantified protein groups. Ellipses indicate 95% confidence regions. (D) Volcano plot of older vs. younger muscle. Dashed lines indicate q = 0.01 and |log2 fold change| = 0.58; 95 proteins were increased (red) and 68 decreased (blue) in older muscle; keratins, hemoglobins, immunoglobulins and other skin- and blood-associated proteins (n = 27) are shown in gray. (E) Top 10 increased and top 10 decreased muscle proteins, excluding skin- and blood-associated proteins, ranked by q-value (left) and by fold change (right), colored by log2 fold change. n = 10 younger (24-30 years) and n = 10 older (70-84 years) women; each data point represents one participant (biological replicate).

### Aging alters the abundance of 163 skeletal muscle proteins

Partial least squares-discriminant analysis (PLS-DA) of all 2,391 quantified protein groups completely separated younger and older participants along the first latent variable (Figure 1C). Differential abundance analysis identified 163 significantly altered protein groups (q < 0.01, |log2 fold change| > 0.58), of which 95 were higher and 68 were lower in older muscle (Figure 1D; Table S1). Twenty-seven of the decreased protein groups were keratins, hemoglobins, immunoglobulins or other skin- and blood-associated proteins, which likely reflect biopsy content rather than muscle biology; these are shown in gray in Figure 1D and were not considered further. Among the most significantly increased proteins were AHNAK, fibrillin-1 (FBN1), tenascin-X (TNXB) and smooth muscle myosin heavy chain (MYH11), while the largest increases among muscle proteins were observed for the metalloproteinase inhibitor TIMP3 (6.6-fold), collagen alpha-1(VIII) chain (COL8A1, 6.2-fold), calponin-1 (CNN1), TPPP3, elastin (ELN) and transgelin (TAGLN) (Figure 1E). Proteins lower in older muscle included glycolytic enzymes (GAPDH, ENO3, LDHA), phosphorylase kinase subunits (PHKA1, PHKB), mitochondrial creatine kinase (CKMT2) and myosin-binding protein H (MYBPH), which showed the largest decrease (7.4-fold) (Figure 1E). Because BMI differed modestly between the age groups, we tested whether BMI could account for the observed age effects. In linear models including age group and BMI as covariates, 110 of the 163 significantly altered proteins retained a significant age-group effect (FDR < 0.05), and no protein showed a significant association with BMI after FDR correction (Table S4), indicating that the age-associated proteome changes were not explained by differences in BMI.

### Multivariate modeling robustly separates younger and older muscle

Stratified five-fold cross-validation of the PLS-DA model confirmed perfect out-of-fold classification of younger and older participants (AUC = 1.00, accuracy 100%) (Figure 2A), and a permutation test with 500 random label assignments indicated that this discrimination was unlikely to arise by chance (p = 0.002) (Figure 2B). Variable importance in projection (VIP) analysis identified 322 proteins with VIP > 1.5, of which 192 were associated with older muscle and 130 with younger muscle (Figure 2C), and 98 of these overlapped with the 163 differentially abundant proteins. The top VIP-ranked proteins higher in older muscle were dominated by nuclear and chromatin-associated proteins, including histone H1.0 (H1-0), several H2B and H4 histones, MATR3, SUN2, macroH2A1, thymopoietin (TMPO), XRCC6 and lamin B2 (LMNB2), together with heterogeneous nuclear ribonucleoproteins (HNRNPA3, HNRNPM, HNRNPA2B1), the ECM proteins COL8A1, COL18A1 and TIMP3, and the lysosomal protease cathepsin Z (CTSZ) (Figure 2D). Conversely, the top VIP-ranked proteins higher in younger muscle were enriched for glycolytic enzymes (GAPDH, PGK1, TPI1, ENO3), glycogen metabolism proteins (PHKA1, PHKB, PHKG1, GBE1), mitochondrial proteins (AFG3L2, SLC25A12, CYCS) and the Z-disc protein filamin C (FLNC) (Figure 2D). Participant-level heatmaps showed that these patterns were consistent across individuals rather than driven by a few outliers, and unsupervised hierarchical clustering of the 136 significantly altered muscle proteins grouped all participants according to age group (Figure S3).

**Figure 2.**
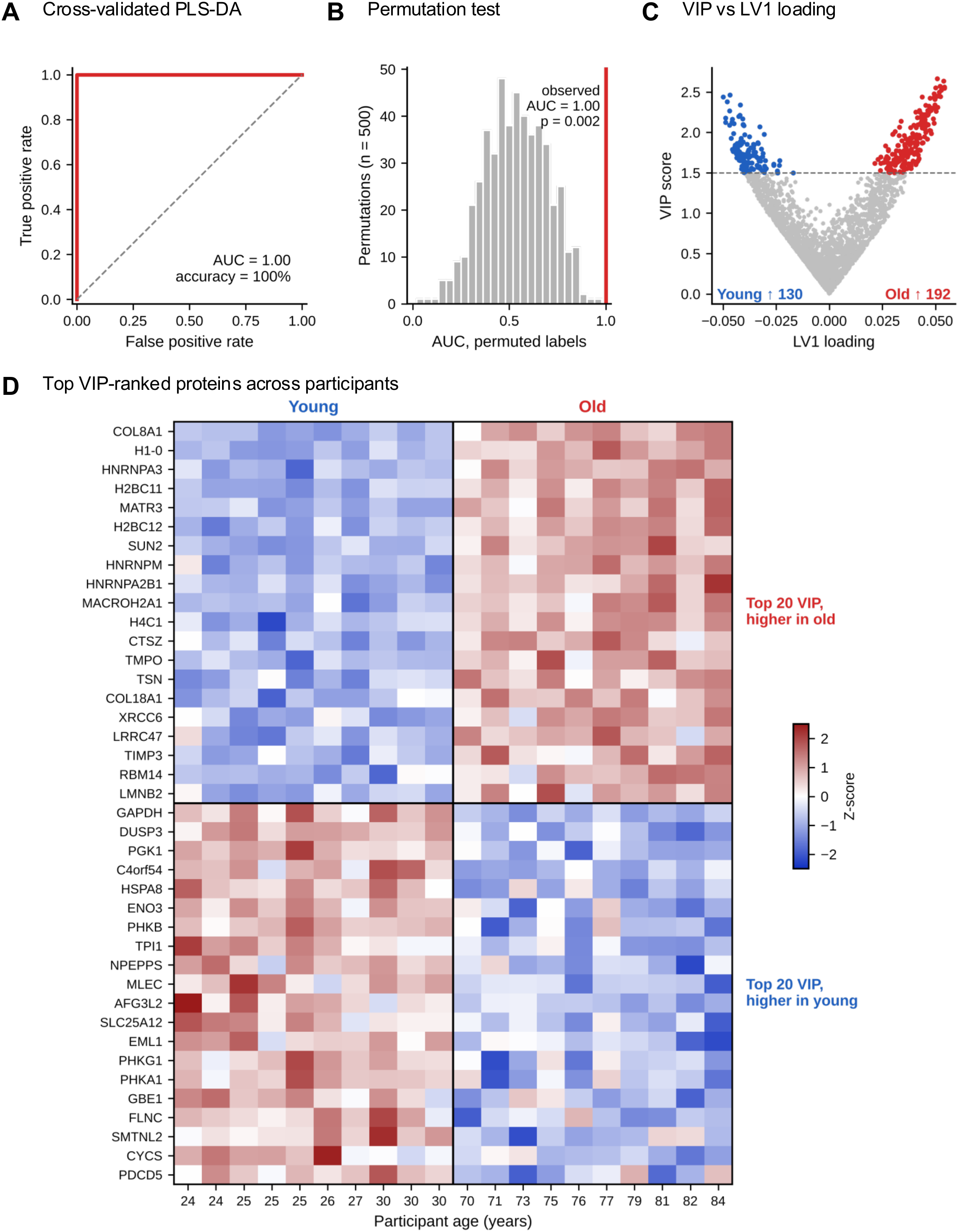
Multivariate modeling robustly separates younger and older muscle. (A) ROC curve from stratified five-fold cross-validated PLS-DA. (B) Distribution of cross-validated AUC values from 500 permutations of group labels; the red line indicates the observed AUC. (C) VIP score vs. LV1 loading for all 2,391 quantified proteins. Proteins with VIP > 1.5 are colored by direction (red, higher in older; blue, higher in younger). (D) Heatmap of the top 20 VIP-ranked proteins associated with each age group. Values are Z-scored log2 intensities; participants are ordered by age. n = 10 younger (24-30 years) and n = 10 older (70-84 years) women; each data point represents one participant (biological replicate).

### Aging shifts muscle toward chromatin, matrix and lysosomal programs at the expense of energy metabolism

Gene Ontology (GO) enrichment analysis showed that proteins higher in older muscle were enriched for chromosome and chromatin organization, chromatin remodeling and assembly, and nucleosome assembly and organization (Figure 3A), whereas proteins higher in younger muscle were enriched for energy derivation by oxidation of organic compounds, ATP metabolic processes, generation of precursor metabolites and energy, and carbohydrate derivative metabolic processes (Figure 3B). Grouping significantly altered proteins by functional annotation highlighted a coordinated shift: chromatin and nucleosome proteins (18 up, 0 down), RNA-binding and spliceosome proteins (12 up, 1 down), structural ECM proteins (13 up, 0 down) and lysosomal proteins (8 up, 1 down) increased with age, whereas glycolysis and glycogen metabolism proteins (0 up, 7 down) and mitochondrial proteins (11 up, 20 down) predominantly decreased (Figure 3C). All 13 significantly altered structural ECM proteins and regulators were increased in older muscle, including TIMP3, COL8A1, ELN, tenascin-C (TNC), CILP, TINAGL1, FBN1, thrombospondin-4 (THBS4), TNXB, periostin (POSTN), COL4A2, biglycan (BGN) and COL18A1 (Figure 3D). Ingenuity Pathway Analysis (IPA) further indicated reduced oxidative phosphorylation and electron transport, altered sirtuin signaling and mitochondrial dysfunction pathways, and changes in creatine-phosphate biosynthesis, glycogen metabolism and glycolysis, together with integrin cell surface interactions and apoptotic execution phase pathways (Figure S4).

**Figure 3.**
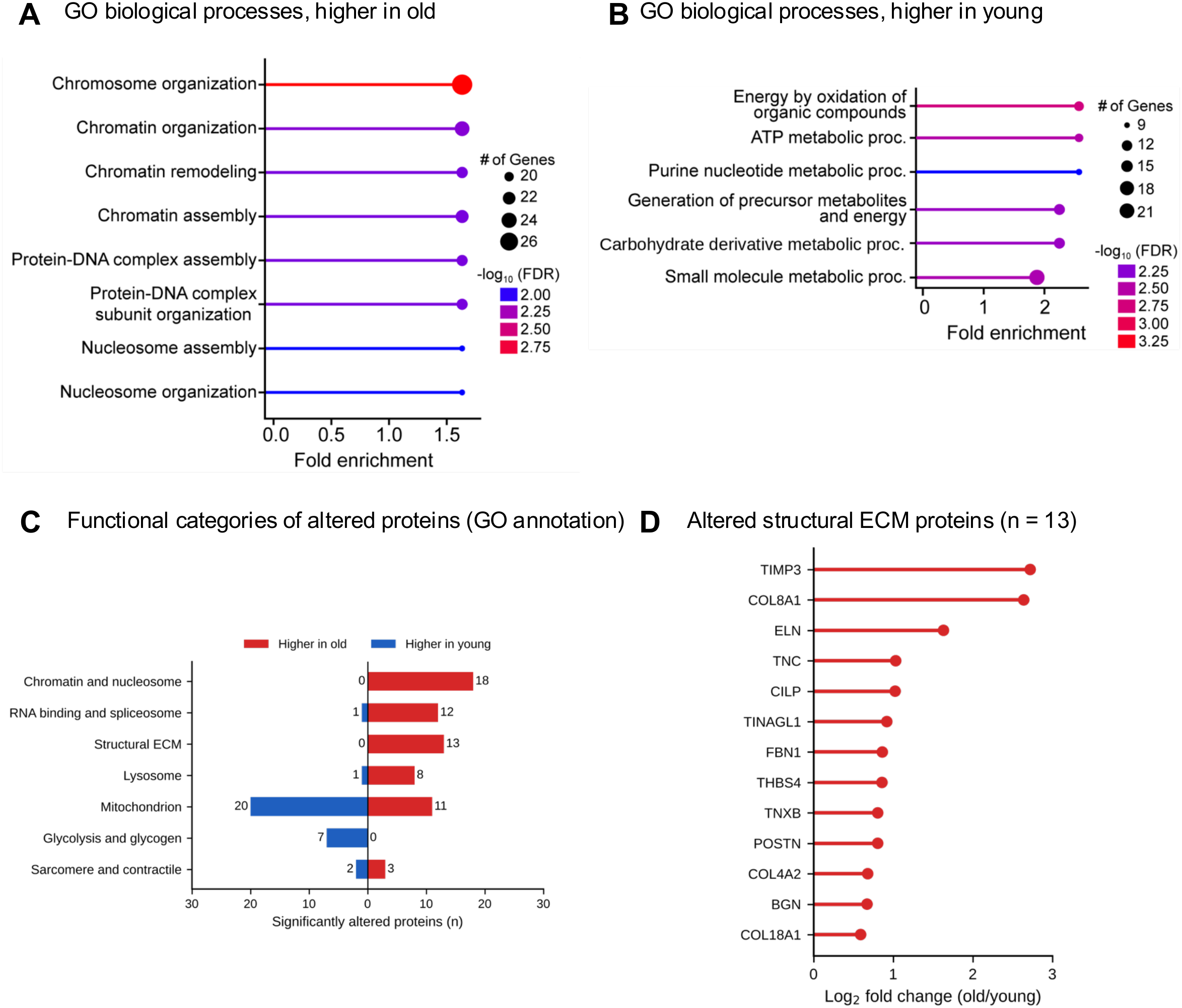
Pathway insights into aging female skeletal muscle. (A, B) GO biological processes enriched among proteins increased (A) and decreased (B) in older muscle. Keratin-driven terms (keratinization, epidermal, keratinocyte and epithelial cell differentiation, intermediate filament organization) are not shown. (C) Numbers of significantly altered proteins increased (red) or decreased (blue) in older muscle within functional categories defined by GO annotation. (D) Log2 fold changes of the 13 significantly altered structural ECM proteins. n = 10 younger (24-30 years) and n = 10 older (70-84 years) women; each data point represents one participant (biological replicate).

### A 24-protein skeletal muscle aging signature

To summarize the age-associated changes into a robust and interpretable signature, we integrated three criteria: significant differential abundance (q < 0.01, |log2 fold change| > 0.58), VIP > 1.5, and consistent direction in all ten older participants relative to the median of the younger group. Of the 163 significantly altered proteins, 98 had VIP > 1.5, and 65 of these changed in the same direction in all ten elderly women. From these 65 candidates, we selected the top VIP-ranked proteins within four biological categories to define a 24-protein skeletal muscle aging signature, termed Buck SENmuscle 1.0 (Figure 4A; Table S5): matrix and vascular remodeling (COL8A1, TIMP3, ELN, FBN1, COL18A1, TNXB); nuclear, chromatin and RNA-binding (H1-0, HMGB1, HMGB2, MATR3, NCL, XRCC6, HNRNPA3, SUN2); lysosomal and SASP-associated (CTSZ, CTSD, PSAP); all increased with age, and glycolytic and energetic proteins decreased with age (GAPDH, ENO3, PHKA1, PHKG1, MYBPH, CYCS, NDUFB4). A participant-level directional signature score, calculated as the mean Z-score of the 17 increased proteins minus the mean Z-score of the 7 decreased proteins, completely separated younger from elderly women (p = 9.5 x 10^-11^) (Figure 4B) and correlated with chronological age (r = 0.96) (Figure 4C). Because ages in this cohort cluster in two groups, this correlation primarily reflects the separation between younger and elderly women rather than a continuous age trajectory.

**Figure 4.**
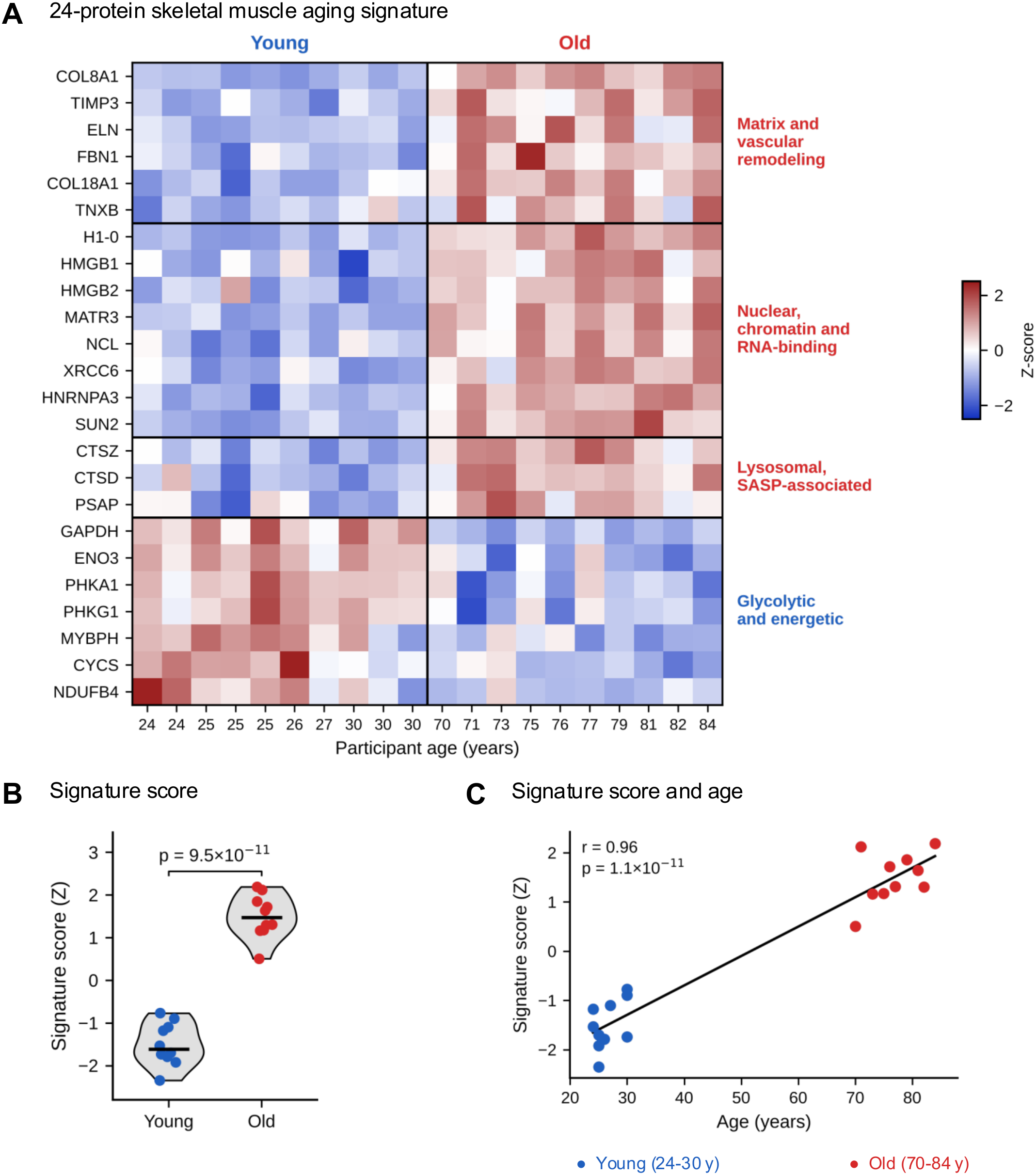
Aging signature development. (A) Heatmap of the 24-protein skeletal muscle aging signature (Buck SENmuscle 1.0) grouped into four biological categories; Z-scored log2 intensities, participants ordered by age. Signature proteins were selected in four steps: (1) significant differential abundance in Spectronaut (q < 0.01 and |log2 fold change| > 0.58; 163 proteins); (2) PLS-DA VIP > 1.5 (98 of 163); (3) same direction of change in all 10 elderly women relative to the median of the younger group (65 of 98); and (4) top VIP-ranked proteins per biological category (24). (B) Participant-level directional signature score, calculated as the mean Z-score of the 17 increased proteins minus the mean Z-score of the 7 decreased proteins (unpaired t-test). (C) Signature score vs. chronological age; the line shows the ordinary least-squares linear regression fit across all 20 participants, with Pearson r and p-value indicated. n = 10 younger (24-30 years) and n = 10 older (70-84 years) women; each data point represents one participant (biological replicate).

### The muscle aging signature tracks a combined measure of physical function

To capture overall physical performance in a single measure, we calculated a combined function score as the mean of Z-scored grip strength and Z-scored gait speed. Grip strength (p = 6.7 x 10^-5^), gait speed (p = 0.020) and the combined function score (p = 3.3 x 10^-5^) were all lower in elderly women (Figure 5A). All 24 signature proteins correlated with the combined function score across participants in the expected direction, with proteins increased with age correlating negatively and proteins decreased with age correlating positively with function (Figure 5B). Twenty-three of the 24 signature proteins correlated with the combined function score at FDR < 0.05, compared with 21 for grip strength and 8 for gait speed alone (Figure 5C; Table S5), indicating that integrating strength and mobility captures the functional relevance of the signature more completely than either measure alone. Because function and protein abundance both differ strongly by age group, we also tested associations within age groups. Most associations were attenuated after adjustment for age group, but HMGB1 (partial r = −0.66, p = 0.002), HMGB2 (partial r = −0.55, p = 0.015) and XRCC6 (partial r = −0.50, p = 0.030) remained associated with the combined function score (Figure 5B). Across the full proteome, 95 proteins correlated with grip strength at FDR < 0.05, whereas no individual protein reached FDR < 0.05 for gait speed (Figure S5A, B; Table S4). After adjustment for age group, 21 proteins remained associated with grip strength at FDR < 0.05; however, many of these within-group associations involved erythrocyte membrane proteins (SLC4A1, EPB41, EPB42, ANK1, ADD1), suggesting a contribution of variable blood content, and these associations should be regarded as exploratory (Figure S5C).

**Figure 5.**
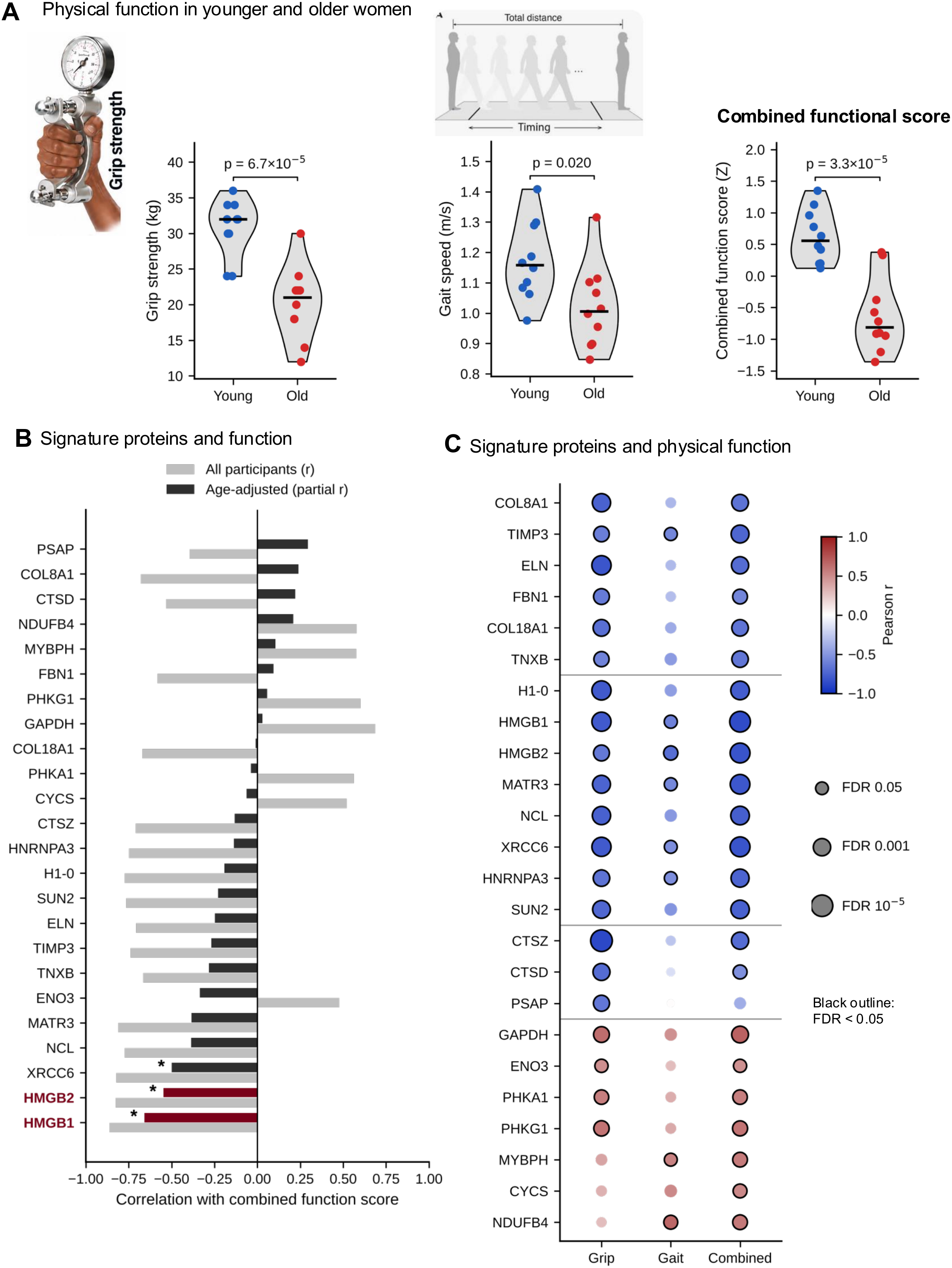
The muscle aging signature tracks physical function. (A) Grip strength, gait speed and combined function score (mean of Z-scored grip strength and Z-scored gait speed) in younger and elderly women (unpaired t-test). Illustrations depict the grip strength (hand dynamometer) and gait speed (timed walk) assessments. (B) Correlation of the 24 signature proteins with the combined function score across all participants (gray) and after adjustment for age group (dark; HMGB1 and HMGB2 highlighted). Asterisks indicate age-adjusted p < 0.05. (C) Correlation of each signature protein with grip strength, gait speed and the combined function score. Color indicates Pearson r, size indicates FDR (Benjamini-Hochberg across the 24 proteins), and black outlines mark FDR < 0.05. n = 10 younger (24-30 years) and n = 10 older (70-84 years) women; each data point represents one participant (biological replicate).

### Signature proteins that indicate functional output

At the participant level, the signature score correlated strongly with the combined function score (r = −0.79, p = 2.9 x 10^-5^) (Figure 6A). The proteins most strongly associated with gait speed were mitochondrial oxidative phosphorylation components that were lower in older muscle, and a composite score of SDHB, NDUFA5, NDUFA8, NDUFB6 and the ADP/ATP translocase SLC25A4 correlated with gait speed (r = 0.76, p = 9.4 x 10^-5^) (Figure 6B); because this score was assembled from the top-ranked proteins, it requires validation in an independent cohort. Among individual signature proteins, the matrix proteins TIMP3 (6.6-fold, q = 3.8 x 10^-4^) and ELN (3.1-fold, q = 1.7 x 10^-7^) showed the largest age-related increases (Figure 6C). However, although both correlated with the combined function score across all participants (TIMP3, r = −0.74; ELN, r = −0.71), these associations were not maintained within age groups (Figure S6B). In contrast, the high mobility group box proteins HMGB1 and HMGB2, chromatin-associated proteins that act as alarmins when released from damaged or senescent cells (Davalos et al., 2013), were higher in all ten older participants (HMGB1, q = 4.6 x 10^-11^; HMGB2, q = 3.6 x 10^-7^) (Figure 6D) and showed the strongest associations with the combined function score of all signature proteins (HMGB1, r = −0.86; HMGB2, r = −0.83) (Figure 6E). Both associations persisted within age groups (HMGB1, age-adjusted r = −0.66; HMGB2, age-adjusted r = −0.55) (Figure 6E), indicating that women with higher muscle HMGB1 and HMGB2 had lower physical function than other women of similar age. Finally, among proteins associated with grip strength, the lysosomal protease cathepsin Z (CTSZ), a component of the signature, showed one of the strongest correlations across the proteome (r = −0.88, FDR = 4.0 x 10^-4^) (Figure 6F), together with LRRC47, AHNAK and ELN (Figure S6A). Figure 7 summarizes the transition from a young muscle proteome characterized by glycolytic, glycogenolytic and oxidative capacity to an older muscle proteome with matrix and vascular remodeling, nuclear and chromatin changes and lysosomal and SASP-associated features, accompanied by lower muscle strength and slower gait.

**Figure 6.**
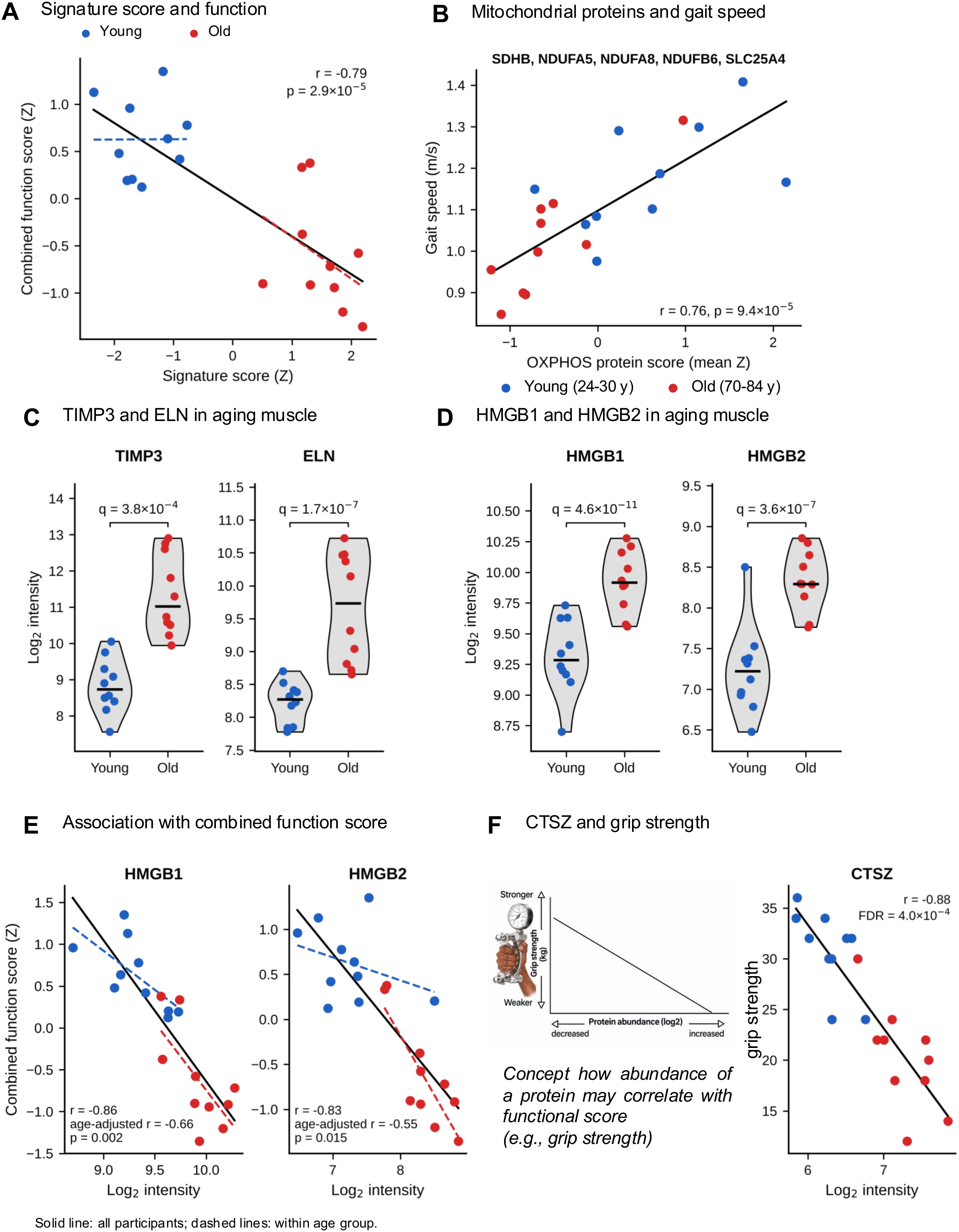
Proteins that indicate functional output. (A) Correlation of the signature score with the combined function score; solid line, all participants; dashed lines, within age groups. (B) Correlation of gait speed with a composite oxidative phosphorylation protein score (mean Z-score of SDHB, NDUFA5, NDUFA8, NDUFB6 and SLC25A4). (C) Log2 intensities of TIMP3 and ELN in younger and older muscle. (D) Log2 intensities of HMGB1 and HMGB2 in younger and older muscle. q-values in (C) and (D) are from Spectronaut. (E) Correlation of HMGB1 and HMGB2 with the combined function score. Solid lines show linear fits across all participants; dashed lines show fits within each age group. Pearson r across all participants and age-adjusted partial r are indicated. (F) Correlation of CTSZ with grip strength (Pearson r; FDR across 2,391 proteins). The inset illustrates the concept of how the abundance of a protein may correlate with a functional score such as grip strength. Blue, younger; red, older participants. n = 10 younger (24-30 years) and n = 10 older (70-84 years) women; each data point represents one participant (biological replicate).

**Figure 7.**
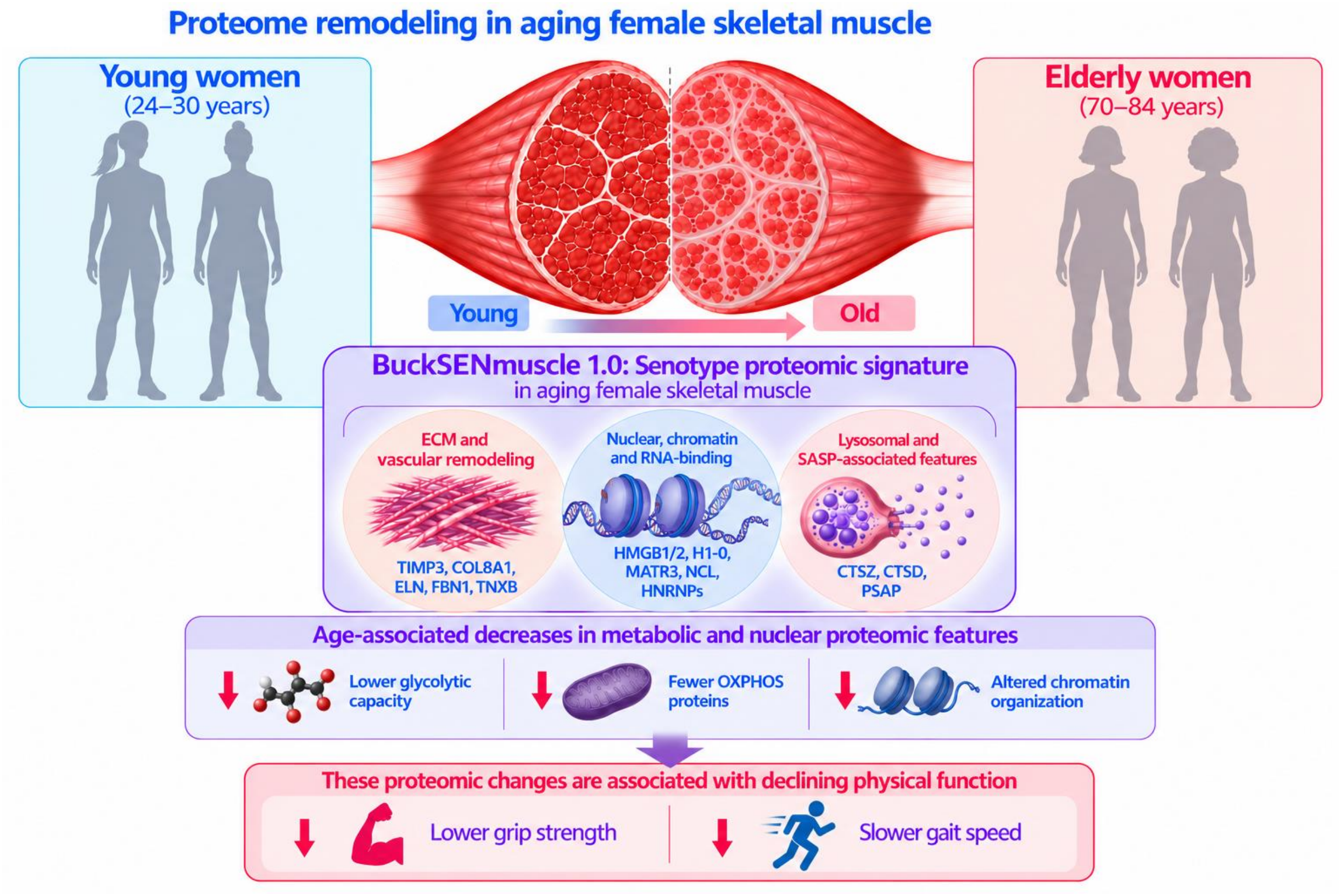
Proteome remodeling in aging female skeletal muscle. Summary of the transition from a younger muscle proteome characterized by glycolytic, glycogenolytic and oxidative capacity, higher grip strength and faster gait speed, to an older muscle proteome with matrix and vascular remodeling, nuclear, chromatin and RNA-binding changes and lysosomal and SASP-associated features, accompanied by lower muscle function. The center of the graphic depicts Buck SENmuscle 1.0, the 24-protein senotype signature, with its three increased components (ECM and vascular remodeling; nuclear, chromatin and RNA-binding; lysosomal and SASP-associated) and age-associated decreases in glycolytic, OXPHOS and chromatin-related features, which are associated with lower grip strength and slower gait speed.

## Discussion

In this study, deep comprehensive DIA proteomic profiling of vastus lateralis biopsies from healthy younger and elderly women revealed a coordinated remodeling of the skeletal muscle proteome with age. Three increased programs, matrix and vascular remodeling, nuclear and chromatin changes, and lysosomal and SASP-associated proteins, together with a decrease in glycolytic and energetic proteins, defined an aging signature that was consistent across individuals and robust to adjustment for BMI. Importantly, this molecular signature tracked physical performance: nearly all signature proteins, and the signature score itself, correlated with a combined measure of grip strength and gait speed, and the nuclear alarmins HMGB1 and HMGB2 remained associated with function within age groups. Our signature also comprises several hallmarks of skeletal muscle health, highlighting that declines in skeletal muscle function may arise from the incorporation of multiple integrated processes. These findings extend previous proteomic studies of aging human muscle to a well-characterized cohort of women and link the proteomic changes directly to clinically meaningful measures of muscle function.

Our results are consistent with the BLSA and GESTALT discovery proteomic findings, which reported lower abundance of energetic proteins and higher abundance of spliceosome and proteostasis proteins in older muscle (Ubaida-Mohien et al., 2019a). Emerging evidence suggests that age-associated alterations in the splicing machinery may reflect dysregulated rather than enhanced splicing activity. In aged mouse skeletal muscle, a markedly attenuated transcript isoform-switching response was described to mechanical overload despite a higher basal splicing score, suggesting that aging may impair the dynamic regulation of RNA processing in response to physiological stress. Together, these findings support broader age-related remodeling of RNA-processing and proteostatic networks in skeletal muscle (Koopmans et al., 2026).

The coordinated decrease of glycolytic enzymes (GAPDH, ENO3, PGK1, TPI1, LDHA), phosphorylase kinase subunits and glycogen branching enzyme in our cohort, together with the decrease of MYBPH, a myosin-binding protein enriched in fast-twitch fibers, is in line with the preferential atrophy of type II fibers with aging and the shift toward a less glycolytic fiber composition (Murgia et al., 2017). Consistent with these findings, Meng et al. also reported age-related declines in glycolytic activity and LDHA expression, which were associated with reduced histone lactylation and increased markers of cellular senescence. These findings suggest that age-related metabolic remodeling in skeletal muscle may extend beyond changes in pathway utilization, potentially influencing epigenetic and transcriptional regulation (Meng et al., 2025). Lower abundance of mitochondrial respiratory chain proteins and mitochondrial creatine kinase is consistent with the decline of muscle oxidative capacity with age, which has been linked to lower walking performance and strength (Zane et al., 2017; Gonzalez-Freire et al., 2018). In our data, a composite of oxidative phosphorylation proteins was associated with gait speed, supporting a link between mitochondrial protein content and mobility in women.

A prominent and unexpected feature of older muscle in this cohort was the increase of nuclear, chromatin and RNA-binding proteins, including linker and core histones, macroH2A1, MATR3, nucleolin and multiple heterogeneous nuclear ribonucleoproteins. Previous research has demonstrated age-associated chromatin remodeling, whereby substantial changes in chromatin architecture and subsequent loss of chromatin loops were shown in muscle stem cells from aged mice. These changes altered interactions between gene promoters and distal regulatory elements and were associated with aberrant gene expression, suggesting that age-related disruption of chromatin organization may contribute to broader changes in transcriptional regulation and, potentially, the proteomic landscape of aged skeletal muscle (Yang et al., 2023).

A similar increase in spliceosomal proteins with age was reported by Ubaida-Mohien and colleagues, who proposed that changes in the splicing machinery may enable muscle cells to respond to accumulating damage (Ubaida-Mohien et al., 2019a). The increase in nuclear proteins could also reflect a relative increase of nuclei from non-myofiber cell types, such as fibroblasts, fibro-adipogenic progenitors, endothelial and immune cells, within aging muscle, or changes in myonuclear chromatin organization. Among these proteins, HMGB1 and HMGB2 are of particular interest in the context of senescence. HMGB1 is an alarmin that is released from senescent cells and promotes SASP signaling (Davalos et al., 2013), and both HMGB1 and the lysosomal cathepsins CTSZ and CTSD were among the proteins most strongly associated with lower grip strength. Furthermore, HMGB1 has been found to negatively regulate skeletal muscle mass (Ho et al., 2022). Most recent, extracellular reduced HMGB1 promotes senescence and SASP-associated signaling in skeletal muscle, whereas inhibition of HMGB1 following muscle injury reduced senescence markers and enhanced muscle regeneration and functional recovery in middle-aged mice (Shin et al., 2025).

In contrast, the canonical senescence marker lamin B1, which is typically lost in senescent cells (Freund et al., 2012), was slightly increased in older muscle (q < 0.01, below the fold-change threshold), and the cell cycle inhibitors p16^INK4a^ and p21^Cip1^ were not detected at the depth of this analysis. These observations indicate that global tissue proteomic workflows capture our new Buck SENmuscle 1.0 senotype signature of the aging muscle environment that likely integrates contributions from senescent and non-senescent cells, consistent with the concept that senescence phenotypes are tissue- and context-specific (Gorgoulis et al., 2019; SenNet Consortium, 2022; Suryadevara et al., 2024).

The increase of structural ECM proteins and regulators, including TIMP3, COL8A1, ELN, FBN1, TNXB, TNC, POSTN, BGN and THBS4, indicates substantial matrix remodeling in aging female muscle, a finding that parallels the fibroinflammatory ECM remodeling we recently described in the postmenopausal ovary (Watson et al., 2026). TIMP3 and COL8A1 showed the largest increases of all quantified proteins, and several ECM proteins, including TIMP3, are components of the SASP (Basisty et al., 2020). Fibrotic matrix deposition increases tissue stiffness and may impair force transmission and regeneration, providing a plausible link between matrix remodeling and the lower grip strength observed in elderly women. Emerging evidence suggests that mitochondrial dysfunction and ECM remodeling may represent an interconnected feature of skeletal muscle aging. This bidirectional crosstalk between these processes whereby mitochondria and oxidative stress can promote profibrotic signaling and ECM accumulation, while ECM deposition and tissue stiffness can further disrupt mitochondrial homeostasis. This could potentially establish a feedforward cycle where tissue fibrosis contributes to the observed aged-related may contribute to decline in gait speed (Cai et al., 2023). We also observed increases in smooth muscle proteins such as CNN1, TAGLN and MYH11, as well as elastin and fibrillin, which together suggest vascular remodeling or a higher relative contribution of vascular structures to the biopsy. Distinguishing between these possibilities will require spatial approaches, such as spatial transcriptomics, MALDI mass spectrometry imaging or other imaging technologies (Gurkar et al., 2023), which are ongoing within our SenNet Buck Tissue Mapping Center.

The 24-protein skeletal muscle aging signature combines significance, multivariate importance and participant-level consistency, an approach analogous to the consensus signature we defined in the human ovary (Watson et al., 2026). The resulting score separated all younger from all older participants and correlated with a combined function score that integrates strength and mobility. Molecular readouts that report on functional performance are of considerable interest for geroscience. Physical function measures such as grip strength and gait speed are robust predictors of disability and mortality but change slowly and are influenced by motivation, pain and other non-muscular factors (Studenski et al., 2014; Cruz-Jentoft et al., 2019; LaMonte et al., 2026). A tissue-level molecular signature that tracks function could complement these measures, help stratify individuals with similar chronological age but different functional trajectories, and provide a mechanistic readout for interventions that aim to preserve muscle function, such as resistance and endurance exercise, nutritional strategies or senotherapeutics (Figure 7). In this context, the persistence of the HMGB1 and HMGB2 associations within age groups is particularly notable, as it suggests that these proteins capture inter-individual differences in functional aging rather than age alone. As HMGB1 is also detectable in circulation, it will be of interest to determine whether muscle-derived signature components can be monitored in plasma, although our matched plasma and tissue analyses indicate that circulating proteins reflect tissue aging only selectively (Huang et al., 2026). As the Buck SENmuscle 1.0 signature was derived and evaluated in the same cohort and the two age groups are widely separated, its performance is likely optimistic. The value of this protein panel as biomarkers correlating wtih muscle function, and its responsiveness to interventions, will need to be tested in independent and longitudinal cohorts that span the full adult age range and include men. Existing human skeletal muscle proteomic data sets, such as those from the Baltimore Longitudinal Study of Aging and the GESTALT study, which span 20 to 87 years of age, include both sexes and were related to physical activity (Ubaida-Mohien et al., 2019a; Ubaida-Mohien et al., 2019b), provide an immediate opportunity to test whether the signature proteins change continuously across adulthood, whether they behave similarly in men, and whether they relate to physical activity.

This study has several limitations. The cohort was small and cross-sectional, and the extreme-group design (24-30 vs. 70-84 years) does not allow modeling of nonlinear trajectories across midlife or the menopausal transition. Only women were included, which strengthens conclusions for female muscle aging but limits generalization to men. The analysis quantified 2,391 protein groups, and low-abundance regulators, including canonical senescence markers, were not detected. Muscle biopsies contain variable amounts of blood and connective tissue, which may contribute to some of the observed differences. ECM categories were defined using GO annotations, and a formal MatrisomeDB analysis (Naba et al., 2012) and overlap with dedicated senescence gene sets will further refine these findings. Finally, we did not perform orthogonal validation by immunohistochemistry in this study. Despite these limitations, the consistency of the changes across participants, their robustness to BMI and their association with grip strength and gait speed support a coordinated, functionally relevant remodeling of the female skeletal muscle proteome with aging.

In conclusion, aging female skeletal muscle shows a shift from a glycolytic and oxidative energetic profile toward matrix and vascular remodeling, nuclear and chromatin changes and lysosomal and SASP-associated features. The resulting senotype 24-protein signature, Buck SENmuscle 1.0, led by the senescence-associated alarmins HMGB1 and HMGB2, links these molecular programs to grip strength, gait speed and their combination, and provides a candidate molecular readout of muscle function for future studies of muscle aging, senescence and interventions that aim to preserve physical performance in women. Importantly, Buck SENmuscle 1.0 connects tissue-level molecular changes with functional metrics in humans that are routinely measured in the clinic, a link that has rarely been demonstrated in human skeletal muscle proteomics.

## Materials and Methods

### Human skeletal muscle biopsy acquisition

Skeletal muscle biopsies were obtained from 20 healthy women enrolled through the NIH Common Fund Cellular Senescence Network (SenNet) Tissue Mapping Center at Wake Forest University under a single Institutional Review Board (IRB)-approved protocol (WF IRB00077384), with written informed consent from all participants. Participants comprised 10 younger women aged 24-30 years (mean 26.6 ± 2.5 years) and 10 elderly women aged 70-84 years (mean 76.8 ± 4.7 years) (Figure S1; Table S1). Participants were eligible for inclusion if they had no contraindications to anticoagulants or lidocaine, a BMI between 18-40 kg/m^2^, free of communicable diseases and major chronic disease, and were not regularly taking medications that indicate treatment for an excluded chronic condition. Exceptions were made for participants receiving stable pharmacological management of hypertension or hypercholesterolemia. Additional inclusion criteria included being between 18 and 35 years of age or 70 years and older, the ability to provide written informed consent, and the absence of medications that could significantly affect cardiovascular function or compromise the safety of the muscle biopsy procedure.

All participants were instructed to fast overnight prior to the muscle biopsy and muscle biopsies were performed between the hours of 8-10am. Skeletal muscle biopsies were obtained using a standardized percutaneous vastus lateralis biopsy procedure (Zamora et al., 2024). Tissue samples were processed immediately after collection, snap-frozen in liquid nitrogen, shipped on dry ice and stored at −80°C until proteomic analysis. Participant characteristics, including age, BMI, self-reported race/ethnicity, smoking status, alcohol use and medical history, were recorded at the study visit. For proteomic analysis, muscle tissue samples (14-50 mg) were used.

### Physical function assessment

Walking performance was assessed using a standardized 400-meter walk test. Participants were instructed to complete 10 laps around a 40-meter course (20 meters out and back) as quickly as possible without running, while maintaining a sustainable pace. Participants were permitted to take standing rest breaks as needed, but the test was discontinued if they were unable to resume walking after 60 seconds or experienced symptoms that compromised their safety. An examiner monitored participants throughout the test and provided standardized encouragement. Total walking time, distance completed, and number of rest breaks were recorded. Heart rate was measured immediately following completion or termination of the test.

Grip strength was measured with a Jamar hydraulic-help dynamometer (JLW Instruments, Chicago, IL). Participants were asked to perform two trails on each hand, and all trials were recorded to the nearest 2 kg.

### Tissue lysis and homogenization

Samples were homogenized in ∼300 µL lysis buffer containing 8 M urea, 2% sodium dodecyl sulfate (SDS), 1 µM trichostatin A (TSA), 3 mM nicotinamide adenine dinucleotide (NAD), 75 mM sodium chloride, and 1X protease and phosphatase inhibitor cocktail (Thermo Fisher Scientific, Waltham, MA) in 200 mM triethylammonium bicarbonate (TEAB) by adding them to 2.0 mL safe-lock tubes (VWR International, Radnor, PA) containing stainless steel beads and subjected to three intervals of high-speed shaking (25 Hz, 1 min) using a Qiagen TissueLyser II (Qiagen, Hilden, Germany). Tissue homogenates were centrifuged at 15,700 x g for 10 min at 4°C, and the supernatant was collected for quantitative proteomic experiments (data-independent acquisitions). Protein concentration was determined using the bicinchoninic acid (BCA) assay (Thermo Fisher Scientific).

### Protein digestion and desalting

Aliquots of 100 µg protein lysate for each sample were brought to the same overall volume of 50 µL with water, reduced using 20 mM dithiothreitol in 50 mM TEAB at 50°C for 10 min, cooled to room temperature (RT) and held at RT for 10 min, and alkylated using 40 mM iodoacetamide in 50 mM TEAB at RT in the dark for 30 min. Samples were acidified with 12% phosphoric acid to obtain a final concentration of 1.2% phosphoric acid. S-Trap buffer, consisting of 90% methanol in 100 mM TEAB at pH ∼7.1, was added, and samples were loaded onto S-Trap mini spin columns. The entire sample volume was spun through the S-Trap mini spin columns at 4,000 x g and RT, binding the proteins to the mini spin columns. Subsequently, S-Trap mini spin columns were washed twice with S-Trap buffer at 4,000 x g and RT and placed into clean elution tubes. Samples were incubated for one hour at 47°C with sequencing-grade trypsin (Promega, San Luis Obispo, CA) dissolved in 50 mM TEAB at a 1:25 (w/w) enzyme:protein ratio. An additional aliquot of trypsin dissolved in 50 mM TEAB was added, and samples were digested overnight at 37°C. Peptides were sequentially eluted from S-Trap micro spin columns with 50 mM TEAB, 0.5% formic acid (FA) in water, and 50% acetonitrile (ACN) in 0.5% FA. After centrifugal evaporation, samples were resuspended in 0.2% FA in water and desalted with Oasis 10-mg Sorbent Cartridges (Waters, Milford, MA). The desalted elutions were subjected to centrifugal evaporation and resuspended in 0.1% FA in water at a final concentration of 1 µg/µL.

### Mass spectrometric analysis

400 ng of each sample were loaded onto EvoTips Pure (Evosep, Odense, Denmark) following the manufacturer’s protocol. During sample loading, one microliter of indexed Retention Time Standard (iRT, Biognosys, Schlieren, Switzerland) was added to each sample. HPLC-MS/MS analyses were performed on an Evosep One liquid chromatography system (Evosep) coupled to a timsTOF HT mass spectrometer (Bruker, Bremen, Germany). The solvent system consisted of 0.1% FA in water (solvent A) and 0.1% FA in ACN (solvent B). Peptides were eluted on a PepSep C18 analytical column (150 µm x 15 cm, 1.5 µm particle size; Bruker) using the 30 SPD method (44-minute gradient length, 500 nL/min flow rate). A zero-dead volume emitter was installed in the nano-electrospray source (CaptiveSpray source, Bruker Daltonics), and the source parameters were set as follows: capillary voltage 1600 V, dry gas 3 L/min, and dry temperature 180°C. Each sample was acquired in data-independent acquisition (DIA) mode coupled to parallel accumulation serial fragmentation (PASEF), or dia-PASEF mode (Meier et al., 2020), with 1 survey TIMS MS scan and 24 dia-PASEF MS/MS scans per 2.03-s cycle. The dual TIMS analyzer was operated at a 100% duty cycle, with equal accumulation and ramp times of 75 ms each. For each TIMS MS scan, the ion mobility ranged from 1/K_0_ = 0.85-1.30 Vs/cm^2^, and the mass range covered m/z 100-1,700. The dia-PASEF scheme was defined as 63 x 15 Th isolation windows from m/z 305.9 to 1,250.9 and covering the ion mobility range 1/K_0_ = 0.74-1.30 Vs/cm^2^. The collision energy was defined as a linear function of mobility, starting from 20 eV at 1/K_0_ = 0.6 Vs/cm^2^ to 59 eV at 1/K_0_ = 1.6 Vs/cm^2^. For calibration of the ion mobility dimension, three ions of Agilent ESI-Low Tuning Mix were selected (m/z [Th], 1/K_0_ [Th]: 622.0289, 0.9915; 922.0097, 1.1986; 1221.9906, 1.13934).

### DIA data processing and statistical analysis

DIA data were processed in Spectronaut v17 (version 17.6.230428.55965) using directDIA (Bruderer et al., 2017; Collins et al., 2017; Gillet et al., 2012). Data extraction parameters were set as dynamic and non-linear iRT calibration, with precision indexed retention time (iRT) selected (Escher et al., 2012). Data were searched against the Homo sapiens reference proteome with 20,423 entries (UniProtKB-TrEMBL), accessed on 01/29/2021. Trypsin/P was set as the digestion enzyme, and two missed cleavages were allowed. Cysteine carbamidomethylation was set as a fixed modification, while methionine oxidation and protein N-terminus acetylation were set as dynamic modifications. Identification was performed using 1% precursor and protein q-value. Quantification was based on the peak areas of extracted ion chromatograms (XICs) of 3-6 MS2 fragment ions, specifically b- and y-ions, with local normalization and q-value sparse data filtering applied. In addition, iRT profiling was selected. Differential protein abundance analysis comparing older (70-84 years) vs. younger (24-30 years) participants was performed using an unpaired t-test, and p-values were corrected for multiple testing using the Storey method (Burger, 2018). Specifically, group-wise testing corrections were applied to obtain q-values. Protein groups with at least two unique peptides, q-value < 0.01, and absolute log2 fold change > 0.58 were considered significantly altered (Table S1). Keratins, hemoglobins, immunoglobulins and other skin- and blood-associated proteins were retained in the statistical analysis but excluded from figure displays and biological interpretation.

### Volcano plot visualization

Differential abundance results were imported from Spectronaut candidate tables containing average log2 fold change, q-values and protein identifiers, and volcano plots were generated in Python. Proteins were classified as increased (log2 fold change > 0.58 and q < 0.01), decreased (log2 fold change < −0.58 and q < 0.01) or not significant.

### Multivariate modeling, VIP analysis and heatmaps

For multivariate analyses, protein intensities were exported from Spectronaut as a pivot table of protein groups, retaining only protein groups supported by at least two unique peptides. Intensities were transformed as log2(intensity + 1), missing values (< 0.1%) were imputed within each protein using the median intensity of that protein across participants, and each protein was standardized to Z-scores before modeling. PLS-DA was performed using a two-component PLS regression framework with age-group labels only (models were not covariate-adjusted). Model performance was evaluated using stratified five-fold cross-validation; within each training fold, features were standardized and a two-component PLS model was fit and applied to held-out samples to generate out-of-fold predictions. Classification performance was summarized using accuracy and receiver operating characteristic (ROC) curves computed from pooled out-of-fold predictions, with the older group as the positive class. Statistical significance of the cross-validated AUC was assessed by repeating the cross-validation procedure with 500 random permutations of the group labels. Variable importance in projection (VIP) scores were computed from the fitted PLS model (Akarachantachote et al., 2014; Chong & Jun, 2005) and plotted against the LV1 loading. Proteins with VIP > 1.5 were considered prioritized and colored by LV1 direction. Heatmaps of the top 20 VIP-ranked proteins associated with each age group were generated using log2-transformed intensities Z-scored across participants, ordered by chronological age.

### Hierarchical clustering

Unsupervised hierarchical clustering of the 136 significantly altered muscle proteins was performed on Z-scored log2 intensities using average linkage and correlation distance for both proteins and participants.

### Pathway and functional category analyses

Gene Ontology biological process enrichment of significantly increased and decreased proteins was performed using Gene Ontology annotations annotated by Spectronaut (tool and background to be confirmed), and canonical pathway analysis was performed using Ingenuity Pathway Analysis (QIAGEN) (Krämer et al., 2014). Functional categories of significantly altered proteins (Figure 3C) were assigned using GO cellular component and biological process annotations exported from Spectronaut. Structural ECM proteins were defined as collagens, glycoproteins, proteoglycans and ECM regulators carrying a GO extracellular matrix annotation.

### BMI sensitivity analysis

To assess whether differences in BMI could account for age effects, each of the 163 significantly altered proteins was modeled by ordinary least squares regression as log2 abundance ∼ age group + BMI. P-values for the age-group and BMI terms were corrected for multiple testing using the Benjamini-Hochberg method (Table S4). This sensitivity analysis complements, and does not replace, the Spectronaut differential abundance analysis.

### Correlation with physical function

Associations between log2 protein abundance and grip strength or gait speed were assessed using Pearson correlation across all 20 participants, with Benjamini-Hochberg correction across all 2,391 proteins (Table S4) or across the 24 signature proteins (Table S5). A combined function score was calculated for each participant as the mean of Z-scored grip strength and Z-scored gait speed. To evaluate associations beyond age group, partial correlations were computed after centering protein abundance and function measures within each age group, with 17 degrees of freedom. An oxidative phosphorylation protein score was calculated as the mean Z-score of SDHB, NDUFA5, NDUFA8, NDUFB6 and SLC25A4.

### Skeletal muscle aging signature

Proteins meeting all three of the following criteria were considered signature candidates: significant differential abundance (q < 0.01, |log2 fold change| > 0.58), VIP > 1.5, and consistent direction in all ten older participants relative to the median abundance of the younger group. From the 65 candidates, the top VIP-ranked proteins within four biological categories were selected to define a 24-protein skeletal muscle aging signature. A participant-level directional signature score was calculated as the mean Z-score of the 17 increased proteins minus the mean Z-score of the 7 decreased proteins.

Python (pandas, NumPy, SciPy, scikit-learn, statsmodels, matplotlib and seaborn) and GraphPad Prism were used for data analysis and visualization.

## Data Availability

The mass spectrometry data and processed data sets supporting the findings of this study are available from the corresponding author upon request.

## Author Contributions

B.S. and J.N.J. conceived and designed the study. H.L.G., J.S., J.N.J. and B.A. recruited participants and collected all muscle biopsies and clinical data. M.A.W. processed samples for mass spectrometry. C.D.K. performed diaPASEF mass spectrometry acquisition. M.A.W. processed diaPASEF data and performed statistical, multivariate and pathway analyses. B.S., M.A.W. and T. R.V. wrote the manuscript. All authors read and approved the final manuscript.

## Acknowledgments

We gratefully acknowledge the women who generously donated skeletal muscle tissue for this study. We acknowledge the commitment and scientific engagement of the late Dr. Judy Campisi and her passionate support of the SenNet program.

## Supplemental Figures

**Figure S1. Participant characteristics.** Age, BMI, self-reported race/ethnicity, smoking status, alcohol use, grip strength, gait speed and tissue amount used for proteomic analysis for each of the 20 participants (10 younger, 10 elderly women).

**Figure S2. Quality control of DIA proteomic data.** (A) Number of protein groups quantified per participant. (B) Coefficients of variation of protein intensities within each age group. (C) Pearson correlation of log2 protein intensities between participants. n = 10 younger (24-30 years) and n = 10 older (70-84 years) women; each data point represents one participant (biological replicate).

**Figure S3. Unsupervised hierarchical clustering of the 136 significantly altered muscle proteins.** Z-scored log2 intensities; average linkage with correlation distance. Skin- and blood-associated proteins were excluded. Note that these proteins were selected based on the age-group comparison. n = 10 younger (24-30 years) and n = 10 older (70-84 years) women; each data point represents one participant (biological replicate).

**Figure S4. Ingenuity canonical pathways.** Canonical pathways identified by Ingenuity Pathway Analysis among significantly altered proteins; color indicates activation z-score and size the gene ratio. The keratinization pathway is not shown. n = 10 younger (24-30 years) and n = 10 older (70-84 years) women; each data point represents one participant (biological replicate).

**Figure S5. Proteome-wide associations with physical function.** (A, B) Pearson correlation of each of the 2,391 quantified proteins with grip strength (A) and gait speed (B) vs. -log10 FDR. (C) Proteins associated with grip strength after adjustment for age group (partial FDR < 0.05). Erythrocyte membrane proteins in this list (SLC4A1, EPB41, EPB42, ANK1, ADD1) may reflect variable blood content of biopsies. n = 10 younger (24-30 years) and n = 10 older (70-84 years) women; each data point represents one participant (biological replicate).

**Figure S6. Additional proteins associated with physical function.** (A) Proteins most strongly correlated with grip strength across all participants (LRRC47, CTSZ, AHNAK, ELN); FDR across 2,391 proteins. (B) Correlation of TIMP3 and ELN with the combined function score; solid lines, all participants; dashed lines, within age groups. Blue, younger; red, older participants. n = 10 younger (24-30 years) and n = 10 older (70-84 years) women; each data point represents one participant (biological replicate).

## Supplemental Tables

**Table S1. Differential protein abundance in older vs. younger female skeletal muscle.** Participant description and Spectronaut differential abundance results (older vs. younger women; unpaired t-test with Storey q-values): all identified protein groups, quantified protein groups with at least two unique peptides (2,391), and the 163 significantly altered protein groups (q < 0.01, |log2 fold change| > 0.58), listed separately for increased and decreased proteins and sorted by fold change and by q-value.

**Table S2. Physical function measurements.** Grip strength and gait speed for each of the 20 participants (10 younger, 10 older women).

**Table S3. Protein quantification across participants.** Spectronaut protein pivot report with protein group intensities for each of the 20 participants, used for multivariate modeling, signature scores and correlation analyses.

**Table S4. Multivariate, clinical association and BMI sensitivity analyses.** PLS-DA VIP scores and LV1 loadings, Pearson and age-adjusted partial correlations with grip strength and gait speed for all 2,391 quantified proteins, BMI-adjusted regression models for the 163 significantly altered proteins, and the 98 significantly altered proteins with VIP > 1.5.

**Table S5. Buck SENmuscle 1.0, the 24-protein skeletal muscle aging signature.** Category, direction of change, fold change, q-value, VIP and consistency for each signature protein; correlations with grip strength, gait speed and the combined function score (all participants and age-adjusted); signature score correlations; and participant-level log2 intensities, Z-scores, combined function scores and signature scores.

