## Supplementary Figure 1 for "Aging Remodels the Female Skeletal Muscle Proteome Toward Matrix, Chromatin and Lysosomal Programs that Track Loss of Physical Function"

### Supplementary Figure S1

#### Figure S1. Participant characteristics

| SenNet ID | Group | Age | BMI | Race/ethnicity | Smoking | Alcohol | Grip (kg) | Gait (m/s) | Tissue (mg) |
|---|---|---|---|---|---|---|---|---|---|
| SN_0196 | Young | 24 | 32.7 | White | Never | Yes | 24 | 1.17 | 20 |
| SN_0214 | Young | 24 | 26.0 | White | Never | Yes | 30 | 1.41 | 50 |
| SN_0120 | Young | 25 | 28.2 | Hispanic/Latino | Never | Yes | 32 | 1.10 | 20 |
| SN_0141 | Young | 25 | 21.2 | White | Never | Yes | 34 | 0.98 | 20 |
| SN_0230 | Young | 25 | 30.4 | White | Never | Yes | 32 | 1.30 | 14 |
| SN_0114 | Young | 26 | 26.4 | Asian | Never | No | 24 | 1.19 | 20 |
| SN_0234 | Young | 27 | 26.6 | White | Former | Yes | 36 | 1.06 | 50 |
| SN_0200 | Young | 30 | 29.2 | White | Never | Yes | 30 | 1.29 | 20 |
| SN_0250 | Young | 30 | 28.7 | White | Never | No | 32 | 1.08 | 50 |
| SN_0254 | Young | 30 | 34.5 | Black | Never | Yes | 34 | 1.15 | 50 |
| SN_0172 | Old | 70 | 19.3 | White | Never | Yes | 22 | 0.90 | 20 |
| SN_0147 | Old | 71 | 21.4 | White | Never | Yes | 22 | 1.00 | 20 |
| SN_0143 | Old | 73 | 20.0 | White | Former | No | 20 | 1.32 | 20 |
| SN_0132 | Old | 75 | 26.0 | White | Former | No | 24 | 1.02 | 18 |
| SN_0133 | Old | 76 | 22.1 | White | Former | Yes | 12 | 1.10 | 16 |
| SN_0130 | Old | 77 | 21.9 | White | Never | Yes | 14 | 1.07 | 20 |
| SN_0148 | Old | 79 | 31.1 | Black | Former | Yes | 18 | 0.89 | 20 |
| SN_0171 | Old | 81 | 22.2 | White | Never | No | 22 | 0.95 | 20 |
| SN_0139 | Old | 82 | 28.4 | Black | Former | Yes | 30 | 1.11 | 20 |
| SN_0152 | Old | 84 | 32.4 | White | Never | No | 18 | 0.85 | 20 |
