## Supplementary figures and images for "Aging Remodels the Female Skeletal Muscle Proteome Toward Matrix, Chromatin and Lysosomal Programs that Track Loss of Physical Function"

### Supplementary Figure 2

#### Supplementary Figure S2

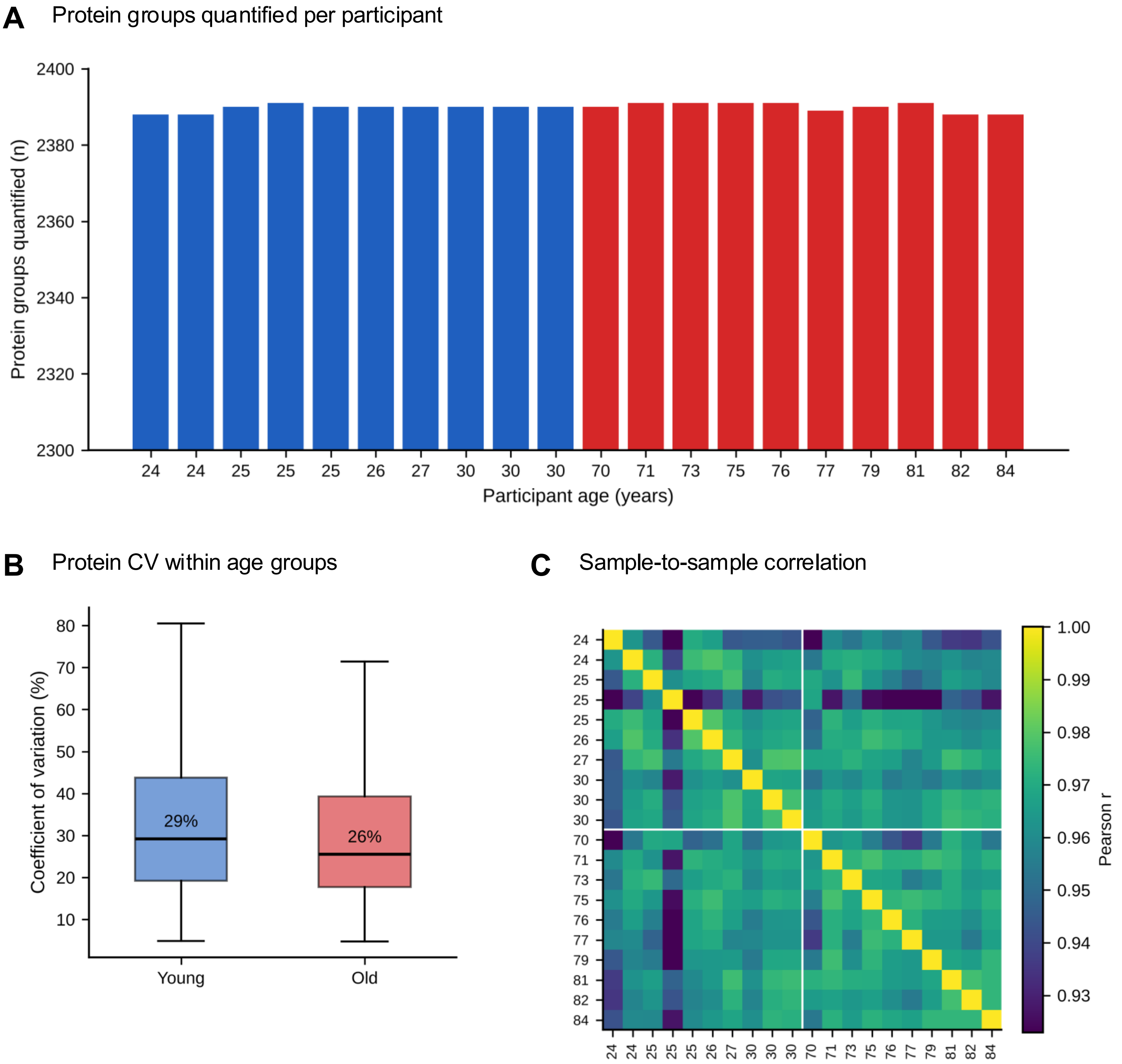

### Supplementary Figure 4

#### Supplementary Figure S4

**A** Ingenuity canonical pathways

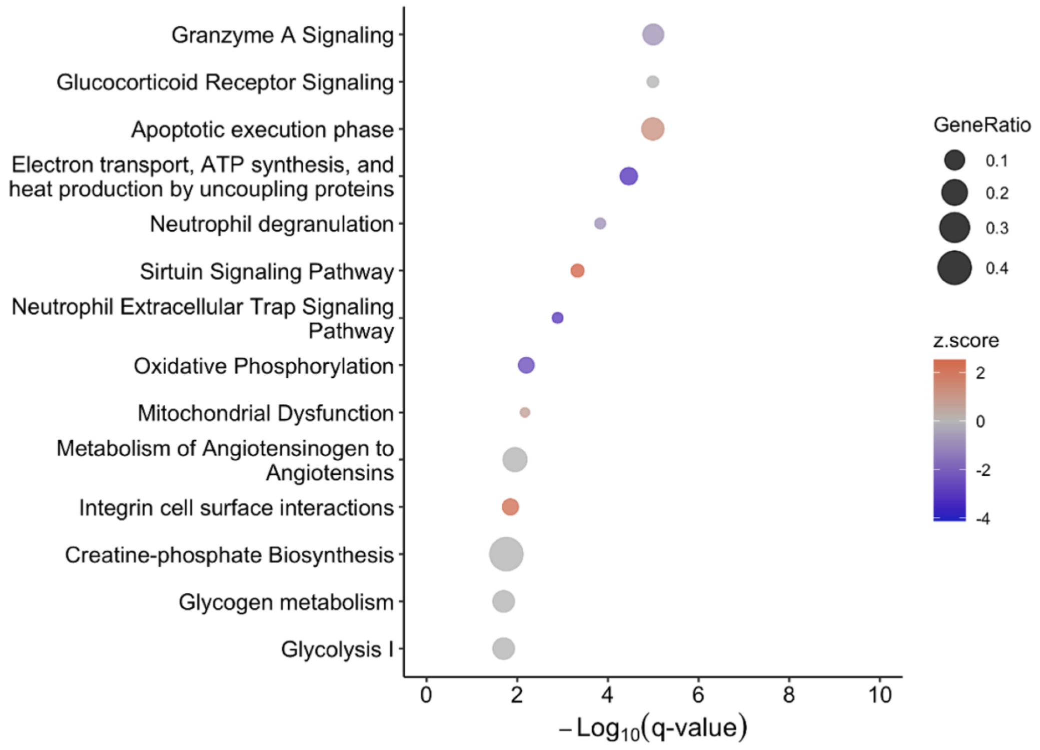
