## Supplementary Figure 3 for "Aging Remodels the Female Skeletal Muscle Proteome Toward Matrix, Chromatin and Lysosomal Programs that Track Loss of Physical Function"

### Supplementary Figure S3

**Figure S3. Unsupervised clustering of 136 significantly altered muscle proteins**

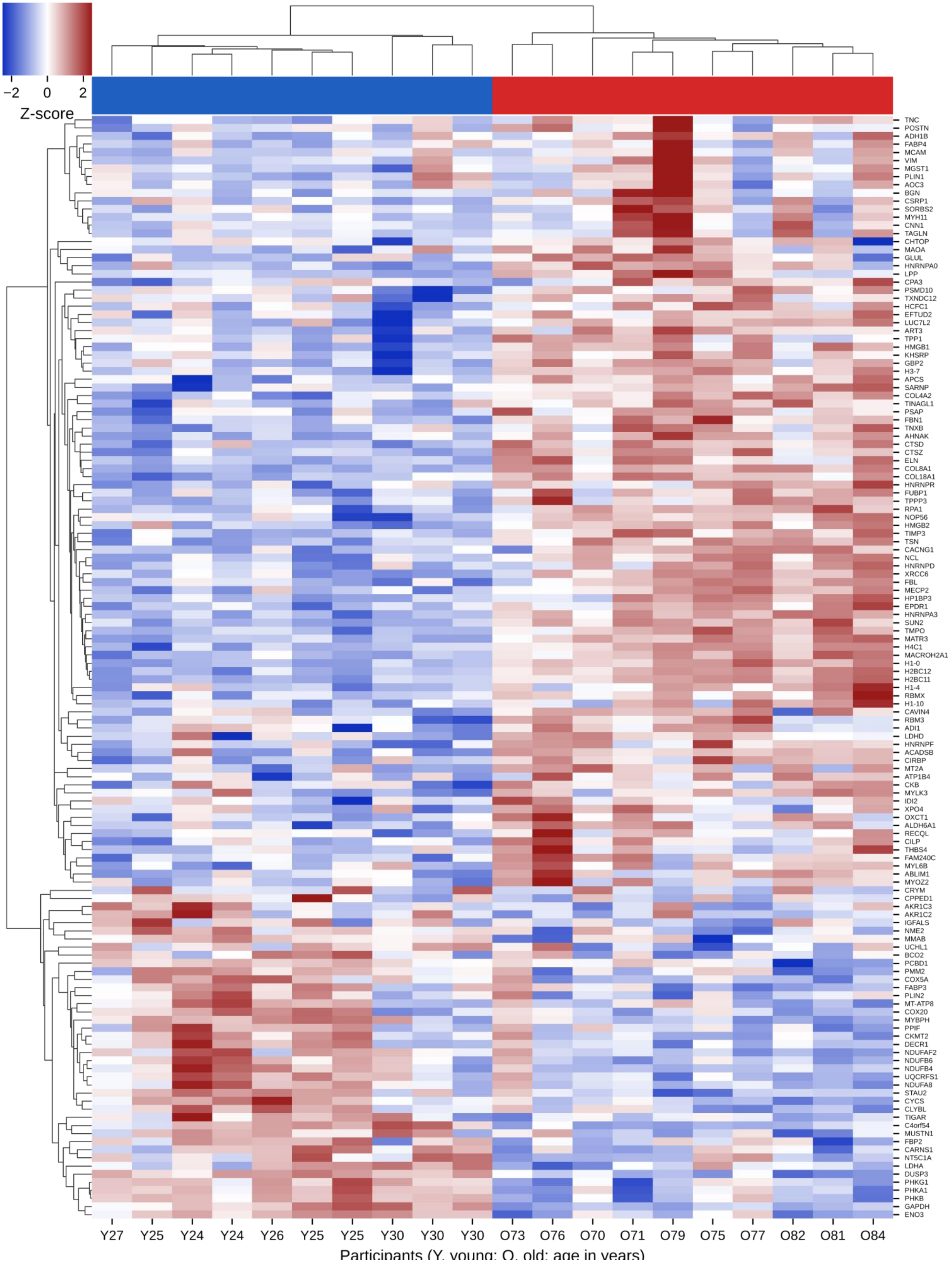
