## Supplementary Figure 5 for "Aging Remodels the Female Skeletal Muscle Proteome Toward Matrix, Chromatin and Lysosomal Programs that Track Loss of Physical Function"

### Supplementary Figure S5

**A** Protein correlation with grip strength

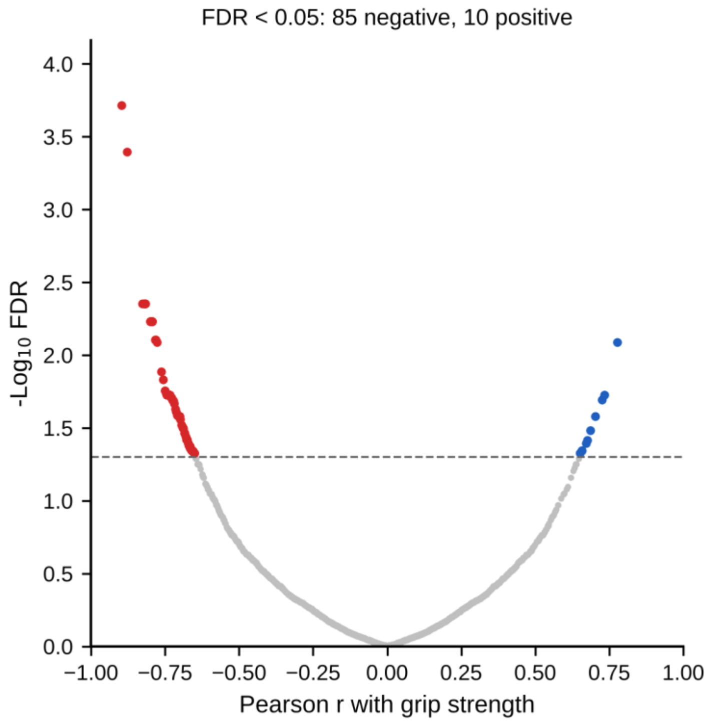


**B** Protein correlation with gait speed

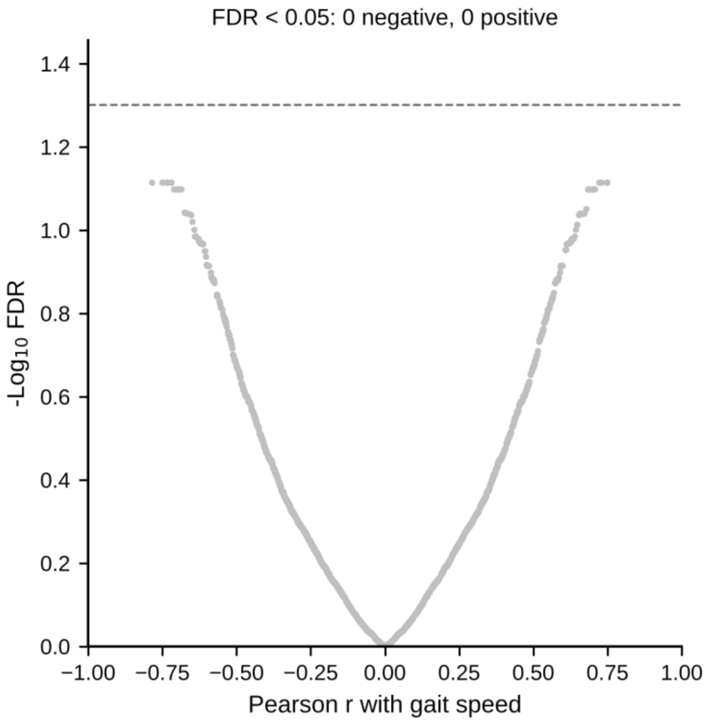


**C** Proteins associated with grip strength after adjustment for age group (n = 21)

| Protein | $Log_2FC$ | q (old/young) | r (grip) | FDR | partial r | partial FDR |
|---|---|---|---|---|---|---|
| IARS1 | -0.36 | 1.5e-03 | 0.06 | 9.1e-01 | -0.83 | 0.014 |
| HSPA5 | -0.04 | 2.4e-01 | -0.34 | 4.3e-01 | -0.81 | 0.014 |
| CHP1 | 0.11 | 3.4e-01 | -0.58 | 1.0e-01 | -0.81 | 0.014 |
| MAPK12 | 0.21 | 1.0e-01 | -0.78 | 7.9e-03 | -0.77 | 0.035 |
| EEF2 | -0.08 | 1.8e-01 | -0.22 | 6.2e-01 | -0.76 | 0.035 |
| TUBB2A | 0.51 | 3.3e-03 | -0.80 | 5.9e-03 | -0.76 | 0.035 |
| EIF1B | -0.07 | 3.2e-01 | -0.42 | 3.0e-01 | -0.76 | 0.035 |
| UFD1 | 0.00 | 5.1e-01 | -0.47 | 2.4e-01 | -0.76 | 0.035 |
| PKLR | -0.07 | 3.5e-01 | 0.52 | 1.8e-01 | 0.75 | 0.039 |
| MAPT | -0.29 | 8.6e-03 | -0.12 | 8.1e-01 | -0.73 | 0.048 |
| SLC4A1 | -0.20 | 2.1e-02 | 0.51 | 1.8e-01 | 0.73 | 0.048 |
| LRRC47 | 0.51 | 5.3e-14 | -0.90 | 1.9e-04 | -0.73 | 0.048 |
| EPB41 | -0.11 | 3.3e-01 | 0.47 | 2.3e-01 | 0.73 | 0.048 |
| MUSTN1 | -0.85 | 3.4e-07 | 0.04 | 9.4e-01 | -0.73 | 0.048 |
| SNCA | 0.06 | 4.2e-01 | 0.43 | 2.8e-01 | 0.72 | 0.049 |
| EPB42 | -0.14 | 2.6e-01 | 0.48 | 2.2e-01 | 0.72 | 0.049 |
| ADD1 | -0.10 | 2.6e-01 | 0.49 | 2.1e-01 | 0.72 | 0.049 |
| RPS25 | 0.03 | 4.4e-01 | -0.51 | 1.9e-01 | -0.71 | 0.049 |
| SORD | -0.42 | 4.2e-02 | 0.73 | 2.0e-02 | 0.71 | 0.049 |
| ANK1 | -0.20 | 3.4e-04 | 0.52 | 1.7e-01 | 0.71 | 0.049 |
| BPHL | -0.52 | 3.1e-02 | -0.21 | 6.6e-01 | -0.71 | 0.049 |
