## Supplementary Figure 6 for "Aging Remodels the Female Skeletal Muscle Proteome Toward Matrix, Chromatin and Lysosomal Programs that Track Loss of Physical Function"

### Supplementary Figure S6

**A** Proteins most strongly correlated with grip strength

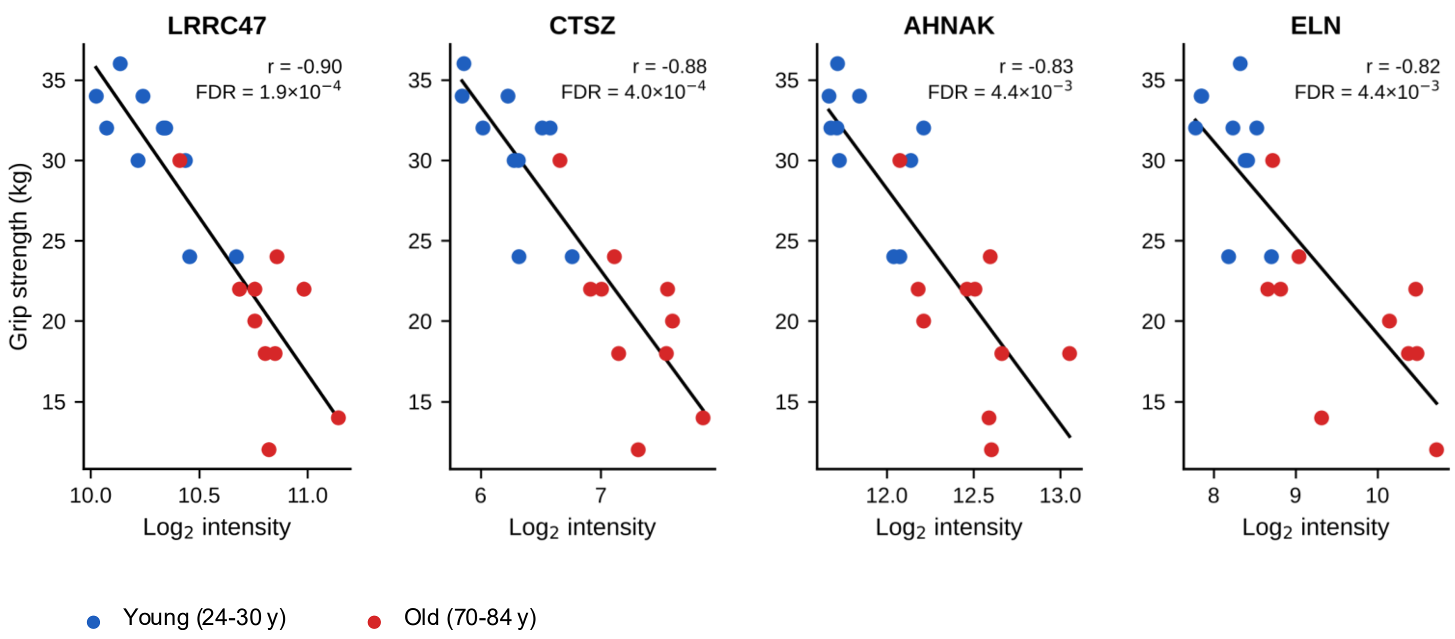


**B** TIMP3 and ELN: association with combined function score

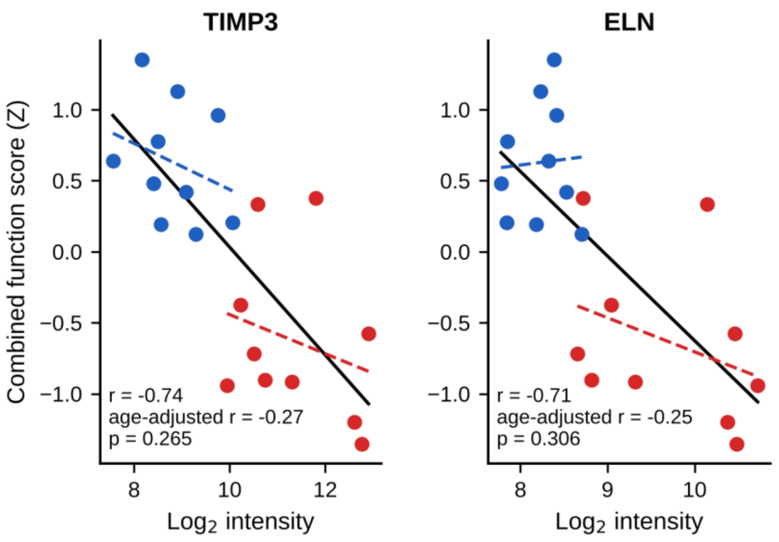


Solid line: all participants; dashed lines: within age group.
